## Supplemental Figures for "*Candida albicans* stimulates the formation of a multi-receptor complex that mediates epithelial cell invasion during oropharyngeal infection"

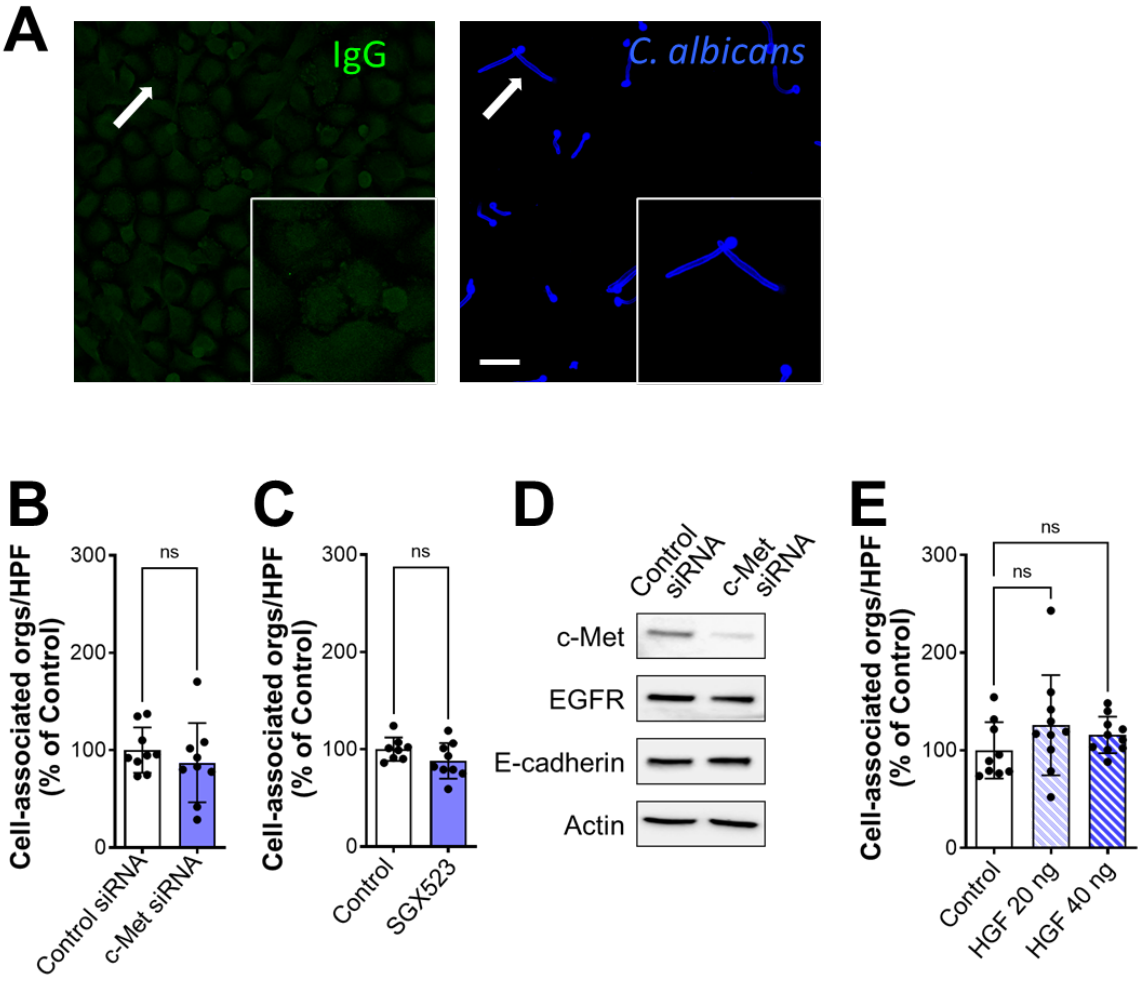


**Fig. S1.** (A) Immunofluorescent image of the OKF6/TERT-2 oral epithelial cell line infected with *C. albicans* and stained with control mouse IgG. Arrows indicate the organism in the magnified inset. Scale bar 10 µm. (B and C) Effects of knockdown of c-Met with siRNA (B) or inhibition of c-Met signaling with SG523 (C) on the number of *C. albicans* cells that were associated (adherent and endocytosed) with the OKF6/TERT-2 oral epithelial cell line. (D) Immunoblot showing the effects of control and c-Met siRNA on the levels of the indicated oral epithelial cell proteins. (E) Effects of hepatocyte growth factor (HGF) treatment on the number of *C. albicans* cells that were associated with oral epithelial cells. Results in (B, C, and E) are mean ± SD of 3 experiments performed in triplicate. ns, not significant (two-way Student's t test [B and C] or one-way ANOVA with Sidak's multiple comparisons test [E])

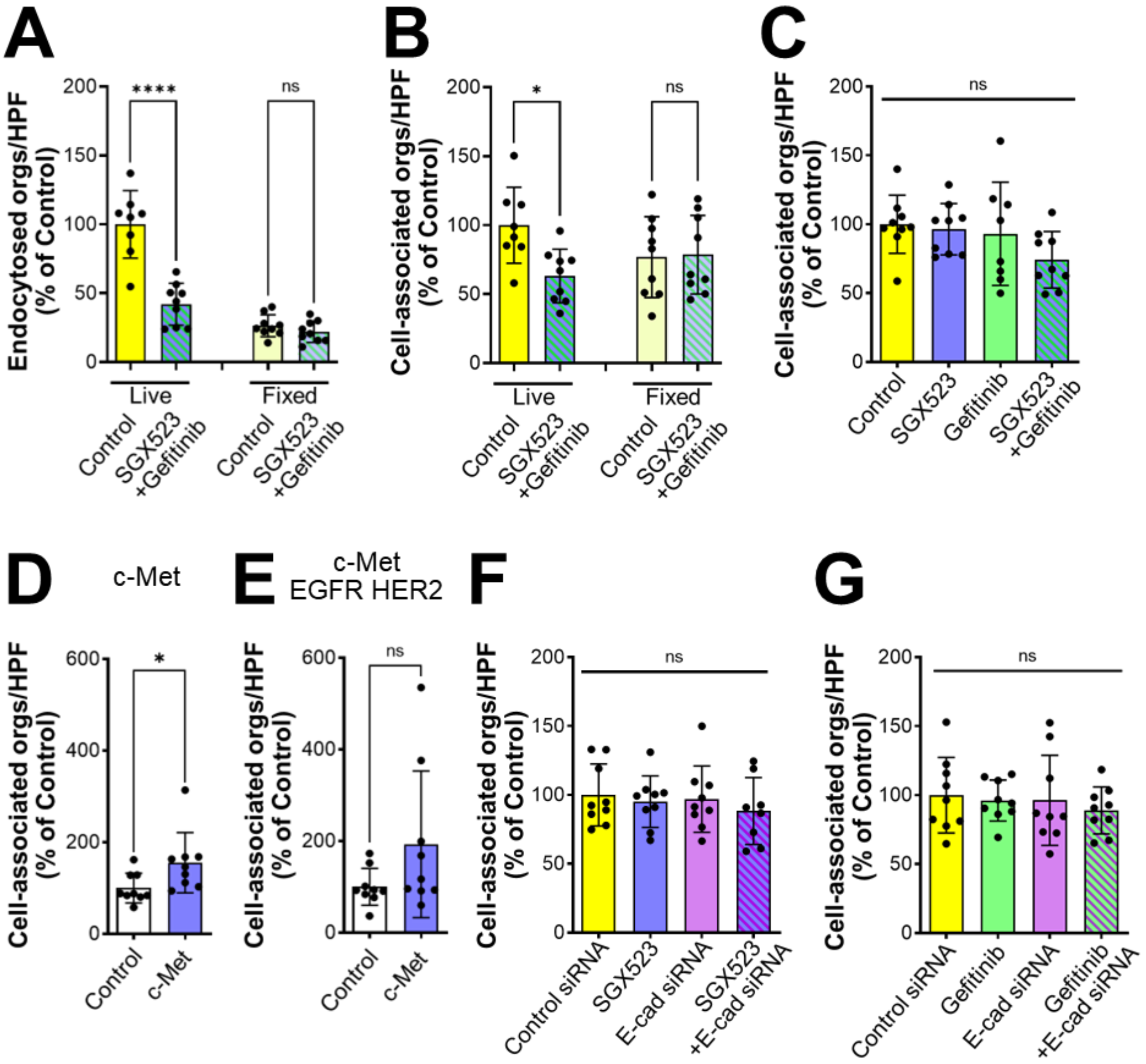


**Fig. S2.** (A and B) Effects of SGX523 and gefitinib on invasion (A) and adherence (B) of *C. albicans* to live and paraformaldehyde-fixed oral epithelial cells. (C) Effects of SGX523 and/or gefitinib on the number of *C. albicans* cells that were associated with oral epithelial cells. (D and E) The number of *C. albicans* cells that were associated with wild-type NIH/3T3 cells (D) or cells expressing the human epidermal growth factor (EGFR) and HER2 (E). (F and G) Effects of siRNA knockdown of E-cadherin in combination with SGX523 (F) or gefitinib (G) on the number of *C. albicans* cells that were associated with oral epithelial cells. Results are mean ± SD of 3 experiments performed in triplicate. *$p$ < 05, **$p$ < 0.01, ***$p$ ,0.001, ****$p$ < 0.0001, ns; not significant (one-way ANOVA with Sidak's multiple comparisons test [A-C, F, G] or two-tailed Student's t test [D and E]).

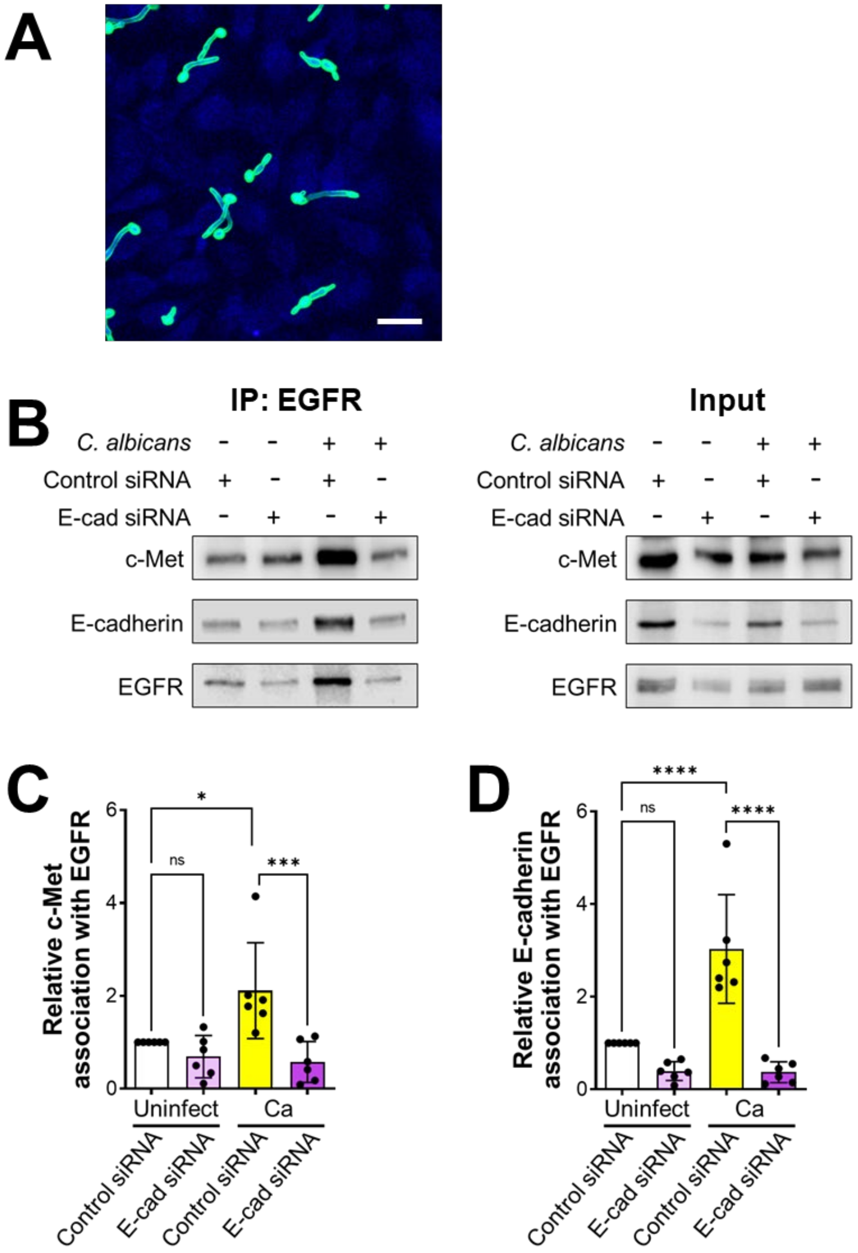


**Fig. S3.** (A) Proximity ligation assay performed with mouse and rabbit IgG as a negative control. Scale bar 10 µm. (B-D) Co-immunoprecipitation experiments in oral epithelial cells transfected with control or E-cadherin siRNA and then infected with *C. albicans* for 20 min. Representative immunoblots of proteins obtained by immunoprecipitation with an anti-EGFR antibody (B). Densitometric analysis of 5 immunoblots (C and D). Results are mean ± SD. **p* < 0.05, ****p* < 0.001, *****p* < 0.0001, ns; not significant (one-way ANOVA with Sidak's multiple comparisons test).

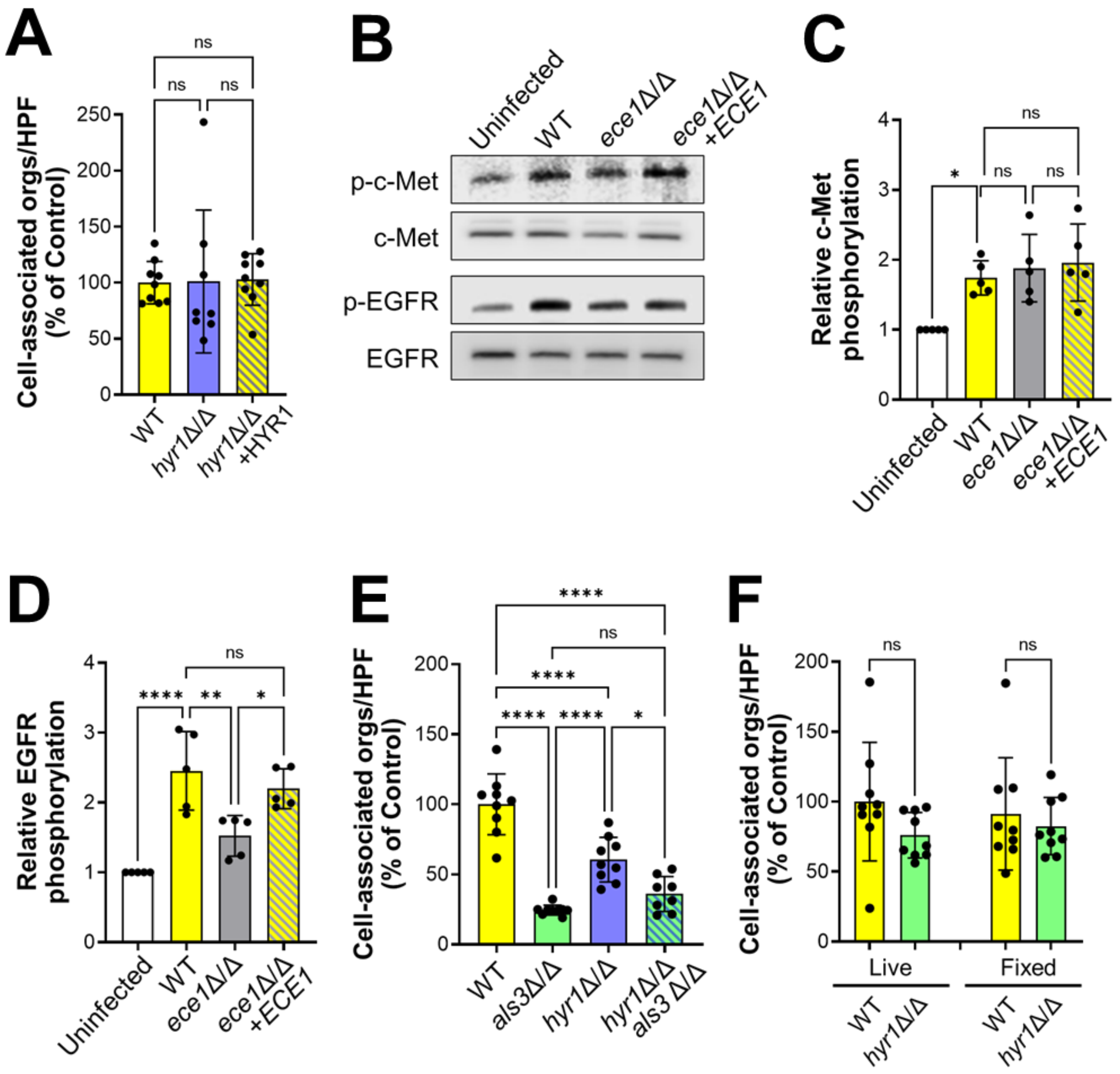


**Fig. S4.** (A) Number of cells of the indicated *C. albicans* strains that were associated with oral epithelial cells. (B-D) Ece1 is dispensable for inducing the phosphorylation of c-Met but not EGFR in oral epithelial cells. Representative immunoblots (B). Densitometric analysis of 5 immunoblots showing the phosphorylation of c-Met (D) and EGFR (E) induced by the indicated strains of *C. albicans*. Results are mean ± SD. (E) Number of cells of the indicated *C. albicans* strains that were associated with oral epithelial cells. (F) Number of cells of the indicated *C. albicans* strains that were associated with live and paraformaldehyde fixed oral epithelial cells.

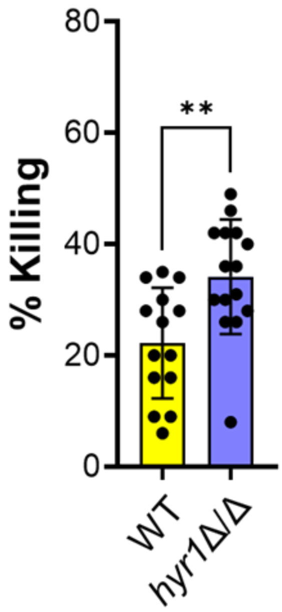


**Fig. S5.** Human neutrophils were infected with the indicated *C. albicans* strains constructed in the SN250 strain background. Results are mean ± SD of neutrophils from 5 donors, tested in triplicate. ***p* < 0.01 (two-sided Student's t test).

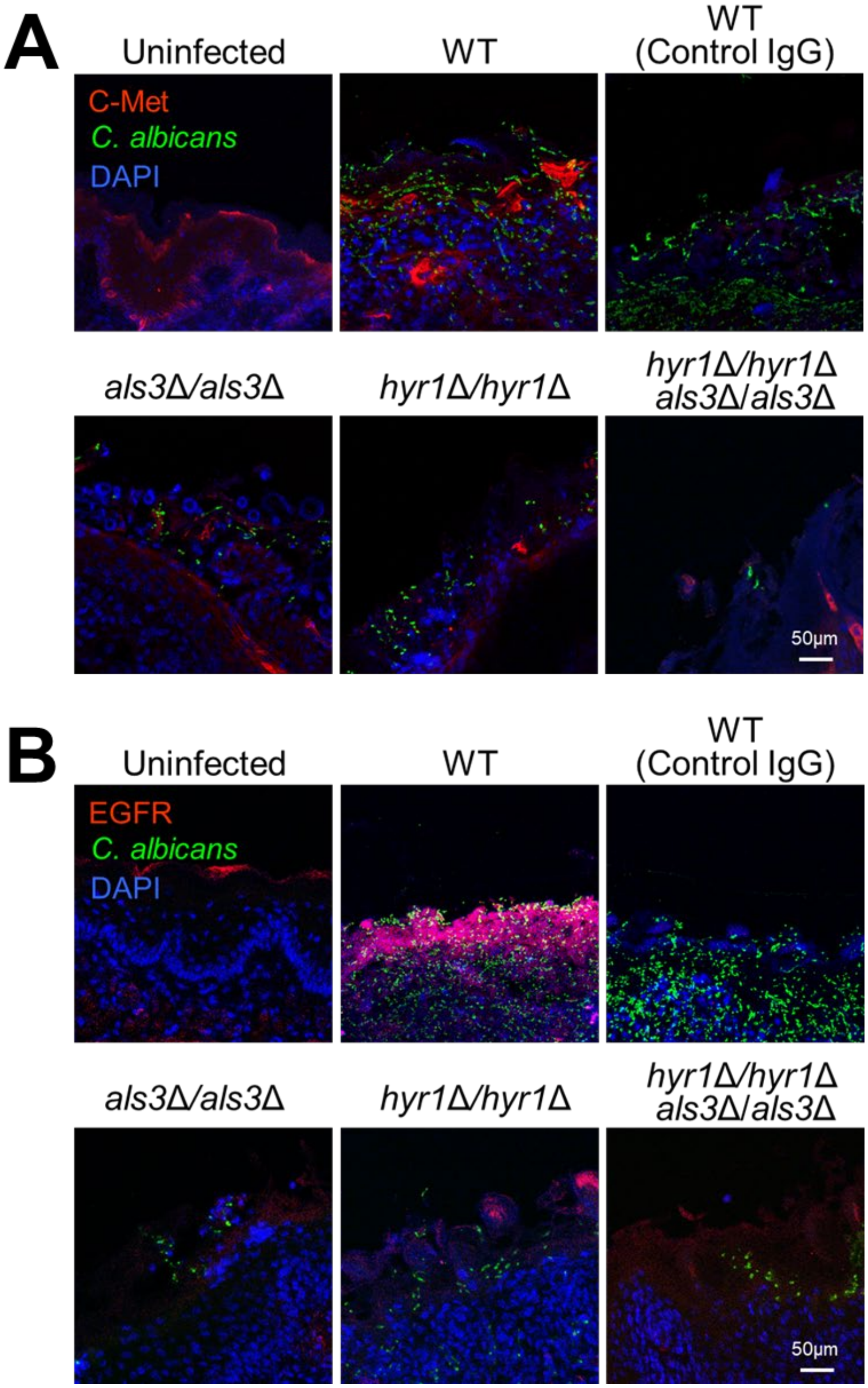


**Fig. S6.** Confocal micrographs of the tongues of immunocompetent mice after 1 d of infection with the indicated strains of *C. albicans*. The samples were stained for c-Met, *C. albicans,* and DAPI (A) or EGFR *C. albicans,* and DAPI (B). Scale bar 50 μm.
