## Supplementary material for "*Candida albicans* stimulates the formation of a multi-receptor complex that mediates epithelial cell invasion during oropharyngeal infection": Table S2

Table S2. List of *Candida albicans* strains used in the work.

| Strain name | Relevant genotype | Reference |
|---|---|---|
| SC5314 | Wild-type | (Jones et al., 2004) |
| SN250 | *URA3/ura3Δ::λimm434 arg4Δ/arg4Δ his1Δ/his1Δ leu2Δ::CdHIS1/leu2Δ::CmLEU2* | (Noble and Johnson, 2005) |
| CAI4 Clp10 | *ura3Δ::λimm434/ura3Δ::λimm434 RPS10/rps10::URA3* | (Brand et al., 2004) |
| MLR63 | *ura3Δ::λimm434/ura3Δ::λimm434 ARG4::URA3::arg4::hisG/arg4::hisG his1::hisG::pTEF1-GFP/his1::hisG* | (Richard et al., 2005) |
| JL036 | *als3Δ::frt/als3Δ::frt* | (Swidergall et al., 2021) |
| MC355 | *hyr1Δ::r1CdHIS1r1, his1Δ::r3* | This work |
| MC374 | *hyr1Δ::r1CdHIS1r1/hyr1Δ::r1CdHIS1r1 his1Δ::r3/his1Δ::r3 als3Δ::r3NAT1r3/als3Δ::r3NAT1r3* | This work |
| MC502 | *hyr1Δ::r1CdHIS1r1/hyr1Δ::r1CdHIS1r1 his1Δ::r3/his1Δ::r3, mdr1Δ::NAT1-HYR1/mdr1Δ::NAT1-HYR1* | This work |
| 823 | *URA3/ura3Δ::λimm434 arg4Δ/arg4Δ his1Δ/his1Δ IRO1/iro1Δ::λimm434 hyr1Δ::CdHIS1/hyr1Δ::CmLEU2* | (Noble et al., 2010) |
| ssa1/als3 | *ura3Δ::λimm434 ssa1::FRT ssa2::FRT als3 rps10::SSA1-URA3 ura3Δ::λimm434 ssa1::FRT SSA2 als3::NAT1 RPS10* | (Liu et al., 2011) |
| JL018 | *ece1Δ::frt/ece1Δ::frt* | (Swidergall et al., 2021) |


Brand, A., MacCallum, D.M., Brown, A.J., Gow, N.A., and Odds, F.C. (2004). Ectopic Expression of *URA3* can influence the virulence phenotypes and proteome of *Candida albicans* but can be overcome by targeted reintegration of *URA3* at the *RPS10* locus. Eukaryot Cell *3*, 900-909.

Jones, T., Federspiel, N.A., Chibana, H., Dungan, J., Kalman, S., Magee, B.B., Newport, G., Thorstenson, Y.R., Agabian, N., Magee, P.T.*, et al.* (2004). The diploid genome sequence of *Candida albicans*. Proc Natl Acad Sci USA *101*, 7329-7334.

Liu, Y., Mittal, R., Solis, N.V., Prasadarao, N.V., and Filler, S.G. (2011). Mechanisms of *Candida albicans* trafficking to the brain. PLoS Pathog *7*, e1002305.

Noble, S.M., French, S., Kohn, L.A., Chen, V., and Johnson, A.D. (2010). Systematic screens of a *Candida albicans* homozygous deletion library decouple morphogenetic switching and pathogenicity. Nat Genet *42*, 590-598.

Noble, S.M., and Johnson, A.D. (2005). Strains and strategies for large-scale gene deletion studies of the diploid human fungal pathogen *Candida albicans*. Eukaryot Cell *4*, 298-309.

Richard, M.L., Nobile, C.J., Bruno, V.M., and Mitchell, A.P. (2005). Candida albicans biofilm-defective mutants. Eukaryot Cell *4*, 1493-1502.

Swidergall, M., Solis, N.V., Millet, N., Huang, M.Y., Lin, J., Phan, Q.T., Lazarus, M.D., Wang, Z., Yeaman, M.R., Mitchell, A.P.*, et al.* (2021). Activation of EphA2-EGFR signaling in oral epithelial cells by *Candida albicans* virulence factors. PLoS Pathog *17*, e1009221.
